# A prospective role for the rumen in generating antibiotic resistance

**DOI:** 10.1101/729681

**Authors:** Cameron R. Strachan, Anna J. Mueller, Mahdi Ghanbari, Viktoria Neubauer, Benjamin Zwirzitz, Sarah Thalguter, Monika Dziecol, Stefanie U. Wetzels, Jürgen Zanghellini, Martin Wagner, Stephan Schmitz-Esser, Evelyne Mann

## Abstract

Antibiotics were a revolutionary discovery of the 20^th^ century, but the ability of bacteria to spread the genetic determinants of resistance via horizontal gene transfer (HGT) has quickly endangered their use^1^. Indeed, there is a global network of microbial gene exchange, the analysis of which has revealed particularly frequent transfer of resistance determinants between farm animals and human-associated bacteria^2^. Here, we leverage the recent release of a rumen microbial genome reference set and show that the wide-spread resistance gene cluster *aadE-sat4-aphA-3* is harboured in ruminal Bacteroidetes. While this cluster appears to have been recently transferred between commensal bacteria in the rumen and many diverse animal and human pathogens, comparative analysis suggests that the cluster stabilized in the pathogens. Then, focusing on streptomycin resistance, it was found that homologues from the rumen span much of the known diversity of aminoglycoside O-nucleotidyltransferases (AadEs) and that distinct variants of the enzyme are present in a single rumen bacterial genome. Notably, a second variant of AadE has also been recently transferred, albeit more often as a single gene, throughout a different set of animal and human associated bacteria. By examining the synteny of AadE orthologues in various bacterial genomes and analyzing corresponding gene trees in an environmental context, we speculate that the ruminant associated microbiome has a salient role in the emergence of specific resistance variants and clusters. In light of the recent literature on the evolutionary origin of antibiotic resistance, we further suggest that the rumen provides a possible route of dissemination of resistance genes from soil resistomes, throughout the farm, and to human pathogens^3^.

---

Since the introduction of antibiotics in 1937, the emergence and spread of antibiotic resistance determinants (ARDs) has become one of the largest threats to human health^1,4^. In fact, the history of antibiotic use is concurrent with the history of increasing antibiotic resistance (AR), which now vastly outpaces new antibiotic discovery^5^. Considering the lack of new antimicrobial compounds entering the clinic, there have been many renewed calls for efforts to discover new compounds or to return to modifying well-understood classes of antibiotics, such as the aminoglycosides^5–8^. In addition to restricting the use of current and future antibiotics, there is a need to better understand the extensive evolutionary history of specific ARDs and their routes of dissemination ^3,9–11^. In doing so, attempting to re-trace evolutionary events involving ARDs and resistance clusters will be essential to move from the metagenomic description of AR reservoirs to identifying particular sources where AR variants emerge, assemble into clusters, and subsequently transfer to human pathogens^12,13^. Fortunately, the ability to carry out such analysis is constantly improving with the number of publicly available genome sequences^14^. Recently, several high-quality datasets containing hundreds of bacterial and archaeal genomes from the rumen microbiome have been published, such as the Hungate1000 collection^15,16^.

In order to search for ARDs that may form clusters in commensal rumen bacteria, we first collected prokaryotic genome sequences from cultured organisms and combined them with a set of metagenome assembled genomes (MAGs), all sourced from the rumen^15,16^. This led to a total of 1585 genomes (453 genomes from cultured organisms and 1133 MAGs). Predicted open reading frames (ORFs) from this dataset were then compared to the comprehensive antibiotic resistance database (CARD) and it was noticed that 2 characterized ARDs from the antibiotic inactivation category, AadE and AphA-3 from *Streptococcus oralis* and *Campylobacter coli*, respectively, each shared 100% amino acid identity with an ORF from three different genomes in the rumen dataset^17^. In all three genomes, these two ORFs were proximal on the same contig, indicating that they may be organized in a cluster (Table S1). Two of the genomes derive from different species of *Bacteroides* from cows in the US, while the third came from a MAG classified as *Prevotella* sampled from a cow in Scotland. When compared at the nucleotide level, the three contigs identified from the rumen bacterial genomes shared a region of approximately ~10kB at 100% nucleotide identity, which upon further annotation, was found to contain the well-known aminoglycoside-streptothricin AR cluster *aadE-sat4-aphA-3*^18^. This cluster was originally identified as the transposon Tn*5405* in *Staphylococcus aureus* and the genes *aadE*, *sat4*, and *aphA-3* encode for an aminoglycoside O-nucleotidyltransferase, a streptothricin N-acetyltransferase and a aminoglycoside O-phophotransferase, respectively (Figure 1)^19,20^. The Tn*5405* sequence itself is also among those conserved at 100% nucleotide identity and to date, the cluster has been observed across a wide range of human and animal pathogens^18–27^. When compared to the NCBI non-redundant nucleotide database, it was found that a highly-conserved region that spanned ~6kB of the ~10kB region was present in a diverse set of pathogens (Figure 1A, Table S2). The segment of this ~6kB which contained the *aadE-sat4-aphA-3* cluster ranged from 99.8-100% nucleotide identity in 32 unique sequences as compared to the rumen sourced contigs, while the flanking regions ranged from 89.9-100% (Figure 1A). Interestingly, the regions that were missing in the pathogens as compared to the rumen bacterial genomes contained only annotated transposases, including a transposase located between *aphA-3* and *sat4* in the cluster, indicating that the cluster has stabilized in the pathogens (Figure 1B, Table S3)^28^. It is worth noting that the only example found where the cluster was not shared as a whole was in *Bacteroides fragilis*, a common reservoir of AR and an opportunistic pathogen, where *aphA-3* appears to have recombined into a different multi-drug resistance cluster, CTnHyb^29,30^. Further, *B. fragilis* was the only non-rumen sequence found with an additional highly conserved region and is the most closely related organism phylogenetically to the three genomes sourced from the rumen. Taken together, the version of *aadE-sat4-aphA-3* identified in rumen *Bacteroides* is highly-conserved in diverse human pathogens, was therefore likely recently horizontally transferred and the loss of transposases, only observed in the pathogenic isolates, implies stabilization of the cluster outside of the rumen. We then sought to gain more evolutionary insight into the individual ARDs within the cluster.

**Figure 1.**
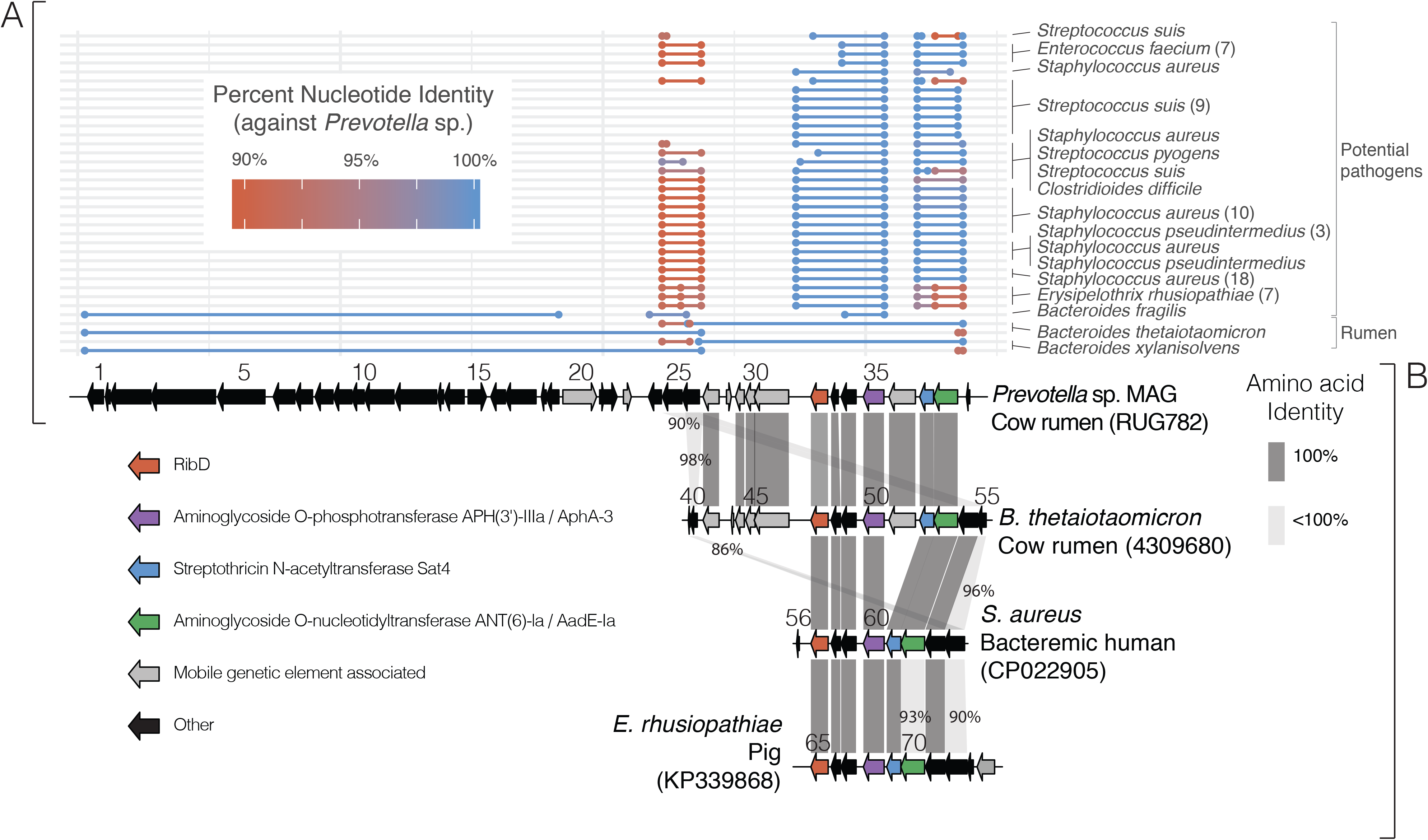
A. Aligned regions from nucleotide blast comparisons of the most similar sequences from NCBI to three rumen contigs (see Table S1) compared to a *Prevotella* sp. metagenome assembled genome (MAG)(RUG782) contig. B. Gene diagram comparison between 2 rumen sourced and 2 pathogen bacterial genomes to show conservation of genetic synteny from a few selected examples. Gene numbering maps to annotations in Table S3 and grey connections between genes represent homologues.

Since genes are the units of evolution and proliferation for mobile traits, we attempted to analyze the evolutionary history of a single enzyme within the cluster. We focused on AadE (also known as ANT(6)), an enzyme characterized to be involved in streptomycin resistance, as it is known to have diverse homologues with the same activity and streptomycin resistance has been long observed in the rumen^31,32^. For instance, in 1966, a range of rumen isolates were screened against various antibiotics and the only compound that demonstrated resistance in all cases was streptomycin^32^. We used 1354 homologues of AadE from the NCBI non-redundant (nr) protein sequences database to build a gene tree (Figure 2)^33,34^. The majority of the homologues (78%) came from Firmicutes, where AadEs likely originated, followed by the Bacteroidetes (13%)^31^. The taxonomic origin of the remaining sequences was diverse and interestingly, despite the fact that only 7% of the sequences derive from the rumen microbiome, they span much of the diversity represented in the NCBI nr database (Figure 2). This indicates that the rumen has been exposed to a large and diverse gene pool with respect to sequences homologous to AadE. Then, we noticed that a sub-clade (clade 7) contained both the AadE from the *aadE-sat4-aphA-3* cluster, as well as a homologous variant from the same rumen *Bacteroides* genome (*Bacteroides thetaiotaomicron* nale-zl-c202 (Hungate collection 4309680)) (Figure 2). These two variants were annotated as ANT(6)-Ia (AadE-Ia) and ANT(6)-Ib (AadE-Ib), respectively. As these two enzymes are thought to have the same activity, we were interested to see how the horizontal transfer of *aadE-Ib* compared with that of *aadE-Ia*^31^. To do so, we carried out the same type of analysis as shown in Figure 1, but instead analyzed the *aadE-Ib* containing contig from the *B. thetaiotaomicron* nale-zl-c202 (Figure 3, Table S2). In this case, *aadE-Ib* was widely distributed in pathogens and commensal bacteria, albeit with lower nucleotide identities as compared to *aadE-Ia* (81.3-100%) and seems to be transferred alone or with a different aminoglycoside O-nucleotidyltransferase (*aad9* or ANT(9)) (Figure 3, Table S3). Considering that the most closely related sequences to *aadE-Ib* are not as conserved and not exclusively found in pathogens, this gene is likely not under as strong of selection as *aadE-Ia*. It is however recombining in context with other ARDs. For example, it was found to recombine near Tet(O) in *C. coli* SX8, a gene which is also highly conserved in several ruminal bacteria at the nucleotide level (Figure 3B, Figure S1A). When looking at further syntenic regions, AadE-Ib was often found in context of AadE-Ia and the *aadE-sat4-aphA-3* cluster. We therefore were interested to further compare AadE-Ia and AadE-Ib across many environments and bacterial genomes and better understand how these two variants may have emerged.

**Figure 2.**
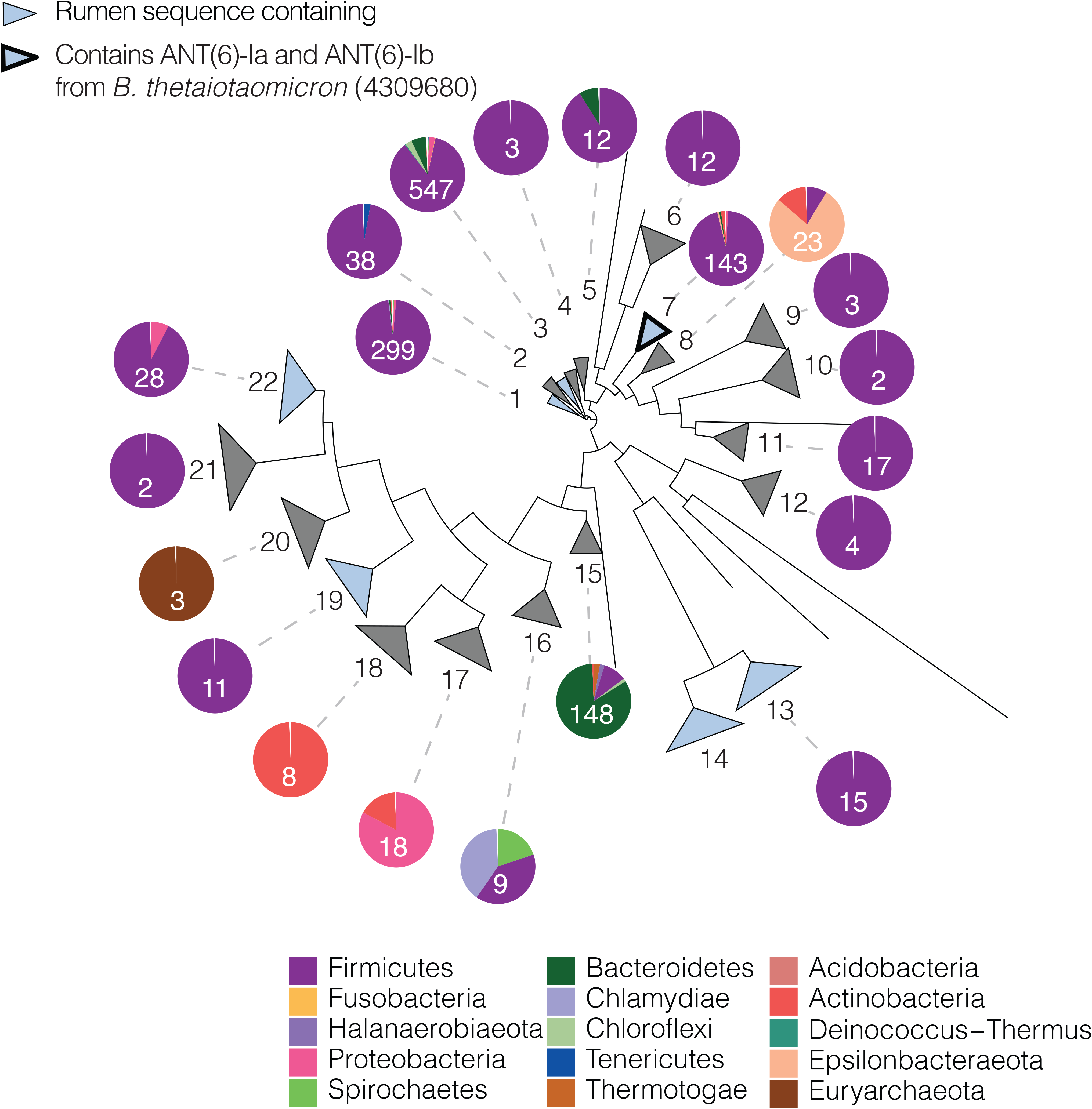
A. Maximum likelihood phylogenetic tree using the top 1354 most similar sequences to AadE-V1 from both the rumen database and NCBI nr with lengths between 250 and 350 amino acids. Clades are numbered for reference. Pie charts show the distribution of phyla from which the sequences were obtained. Numbers within the pie charts indicate how many sequences make up the clade. Clade 7 contains ANT(6)-Ia (AadE-Ia) and ANT(6)-Ib (AadE-Ib) from *B. thetaiotaomicron* nale-zl-c202.

**Figure 3.**
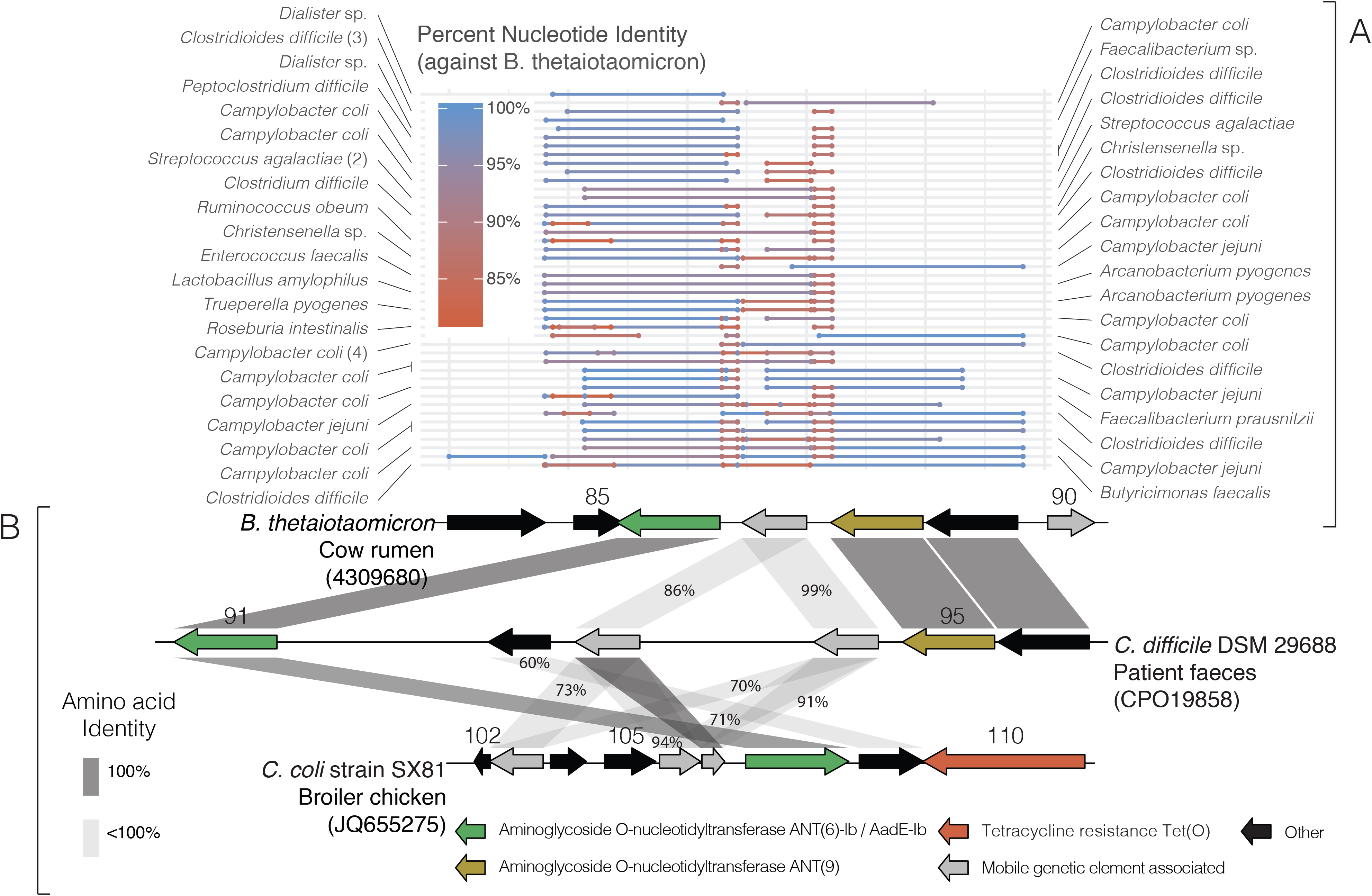
A. Aligned regions from nucleotide blast comparisons of the most similar sequences from NCBI to a single rumen contig (see Table S1) compared to a *B. thetaiotaomicron* nale-zl-c202 genome (Hungate collection 4309680) contig. B. Gene diagram comparison between a rumen sourced and 2 pathogenic organism genomes to show conservation of genetic synteny from a few selected examples. Gene numbering maps to annotations in Table S3 and grey connections between genes represent homologues.

By building a gene tree with all protein sequences within clade 7, shown in Figure 2, we observed four clear sub groups that each corresponded to a different annotated version of AadE (Figure 4A). Outside of bacteria from the rumen or pathogens, the two groups representing AadE-Ia and AadE-Ib contained sequences that were mostly sourced from various animal or human intestinal samples (Figure 4A). Moreover, the sequences from the rumen tended to span these two groups, whereas the sequences from the two more deeply branching sub groups, containing ANT(6)-Id (AadE-Id) and ANT(6)-Ic (AadE-Ic), were mostly sourced from diverse environmental samples, such as plants and soil (Figure 4A, Table S4). This may not be surprising in light of several genomic analyses of the transfer of horizontal resistance, which have pointed to the gut as an interconnection between soil and clinical pathogens or found that farm animal microbiomes are enriched for transfer events with human-associated bacteria^2,11^. This does however point more specifically to the rumen as a link between the environment and the human or animal intestinal tract. Two questions that then arise are: why did these two variants, AadE-Ia and AadE-Ib, emerge evolutionarily and why are they both often present in a single genome (e.g. *B. thetaiotaomicron* nale-zl-c202)? Especially considering that the characterized versions have the same activity^31^. When looking at those genomes that were selected for sharing high nucleotide identity with the *aadE-Ia* or *aadE-Ib* from *B. thetaiotaomicron* nale-zl-c202, several of them were also found to have both or multiple copies of AadE (Figure 4B and Figure 4C). By comparing their identity and synteny, it is clear that several AadEs have arisen via gene duplication events and often remained in context of each other (Figure 4B and Figure C). Interestingly, outside of the *aadE-sat4-aphA-3*, the genes found in context of the two AadEs are mostly streptomycin or aminoglycoside modifying enzymes (Figure 4B and Figure C). Other ARDs in context include tetracycline and lincosidamide resistance genes, which are also heavily represented in the rumen and act on compounds produced by *Streptomyces* (Table S1, Table S5, Figure S1). It is interesting to note, although often observed with other ARDs, that AadE-Ia and AadE-Ib further recombine into clusters with genes which would theoretically yield the same resistance phenotype. A logical suggestion is that aminoglycoside producing bacteria from soil are also the sources of AR, and that these genes may have served modifying roles outside of resistance to the toxicity of the compounds^35–37^. Altogether, it is possible that recombining variants of *aadE* from the environment further duplicated, potentially including the events that spawned AadE-Ia and AadE-Ib, adapted, and refined their syntenic context in the rumen. During the process, there were likely many subsequent transfer events, often with commensal bacteria of the intestinal tract of humans and other animals.

**Figure 4.**
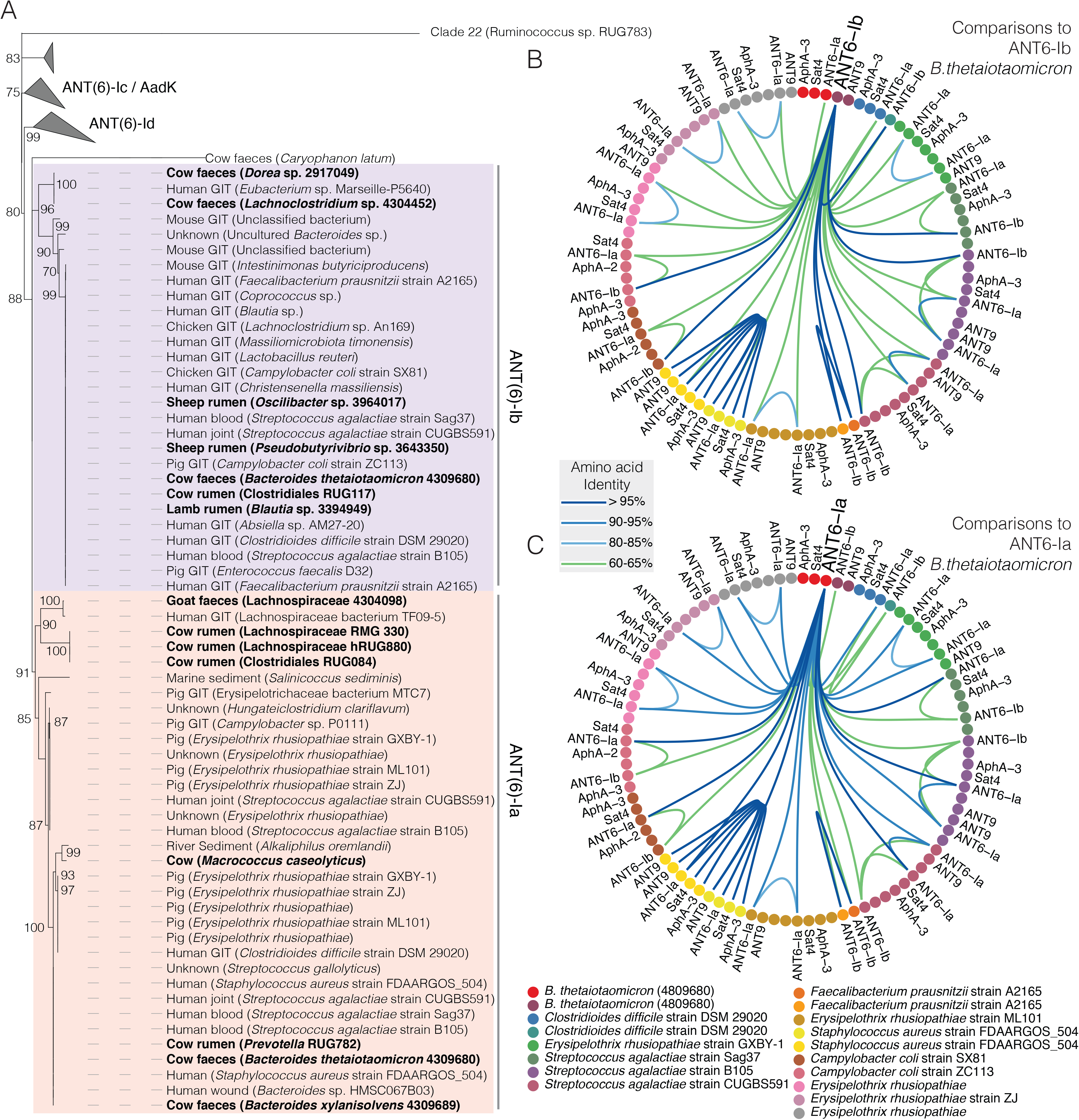
A. A maximum likelihood tree using all sequences falling within clade 7 (Figure 2). Ultrafast bootstrap values are shown and sequences in bold are from the Hungate 1000 collection. Clades are labelled based on containing specific variants of AadE. The outgroup used was a randomly selected sequence taken from clade 22 in Figure 2. B and C. Each point around the circle is an antibiotic resistance determined (ARD) coloured by the contig containing it. A contig is representated if the genome was used in Figure 1 or Figure 3 and contained two or more AadE. An ARD is shown if it is annotated as AadE-Ia or AadE-Ib or another annotated ARD that is syntenic with one of the AadE variants (within a resistance cluster). B. Connections show amino acid identity with AadE-Ib from *B. thetaiotaomicron* nale-zl-c202 (Hungate collection 4309680). C. Connections show amino acid identity with AadE-Ia from *B. thetaiotaomicron* nale-zl-c202 (Hungate collection 4309680). Genes are labelled if they are annotated as AadE-Ib, AadE-Ia, part of the *aadE-sat4-aphA-3* cluster or annotated to act on aminoglycosides. Other ARDs present in the resistance cassettes are shown in Table S5.

In terms of food-producing animals, aminoglycosides accounted for 3.5% of the total sales of antimicrobials in 2015 and are most frequently used to treat infections^38^. Considering the diversity of homologues of aminoglycoside inactivating or modifying enzymes and that cattle are not directly fed aminoglycosides, it is worth considering that the rumen is also exposed to the compounds and an ARD gene pool via natural sources. Soil, for example, is a well characterized reservoir of antibiotic producing organisms and ARDs, which long predate the use of antibiotics, and aminoglycosides have particularly high sorption in soils^8,9,39–41^. Additionally, *Streptomyces* are often isolated from agricultural soils, including in the case of the discovery to streptomycin^42^, as well as from feed sources such as hay directly^43^. The rumen takes in enormous amount of feed and in various ways, it has been shown to provide favourable conditions for genetic exchange^44–46^. Considering that ecology shapes gene exchange, it is reasonable to assume that the rumen, a 100-200L anaerobic bioreactor constantly interfacing with the feed containing a diversity of antibiotic related compounds and the microorganisms producing them, provides an opportunity for a ARD gene pool to exchange and adapt within an animal associated microbiome and environment. While streptomycin is not regularly detected in feed, other compounds produced by *Streptomyces*, which are easier to detect, such as chloramphenicol, are found regularly^47^. Ultimately, a wide range of aminoglycoside modifying enzymes sourced from soils or sediments may be transferred to and refined the rumen, especially in terms of genetic synteny, before being spread throughout the farm and potentially strongly selected or co-selected for when treating an animal infection or when a field is contaminated with antibiotics (Figure S2)^48^. In terms of spreading throughout the farm, the humans, whose associated microbes show 25 fold more HGT as compared to non-human isolates, in contact with the animals are the most obvious conduit^2^. It was however also interesting to find a common dog pathogen (*Staphylococcus pseudointermedius*) in the analysis which contained the highly conserved *aadE-sat4-aphA-3* cluster (Figure 1A). Overall, we observed recent horizontal transfer events of ARDs between ruminal bacteria, farm animals, pets and pathogens infecting humans, whose history of assembly points towards the rumen as the source. Therefore, while only one of many sources of AR, the rumen should be considered an environment with high potential for generating clusters of ARDs and providing a central link to other reservoirs, especially on the farm, before going on to create problems in the clinic. If further evidence corroborates this suggestion, antibiotic discovery efforts could focus on antibiotic compounds from organisms that evolved in environments with little or no connection to agricultural feed.

**Figure S1.**
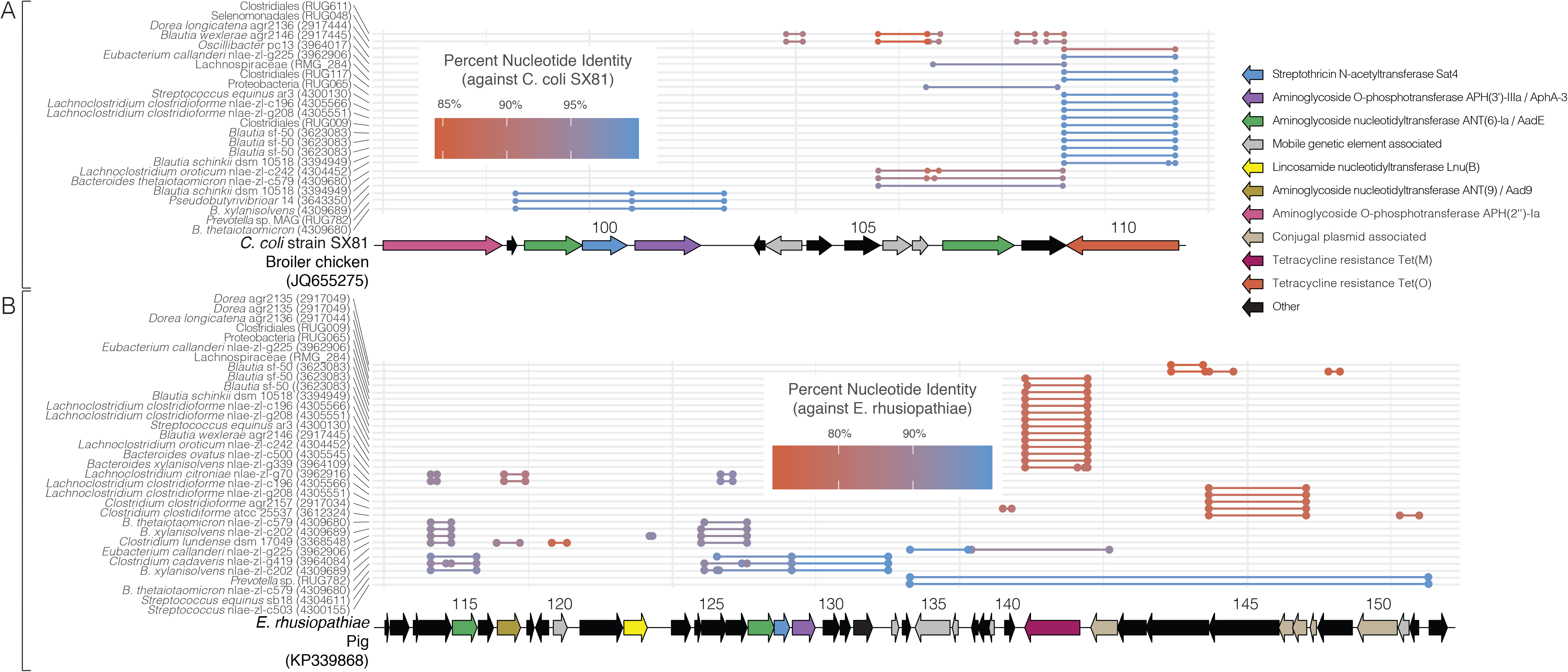
A. Aligned regions from nucleotide blast comparisons of rumen bacterial genomes to a *C. coli* (JQ655275)(A) and *Erysipelothrix rhusiopathiae* (KP339868)(B) genome.

**Figure S2.**
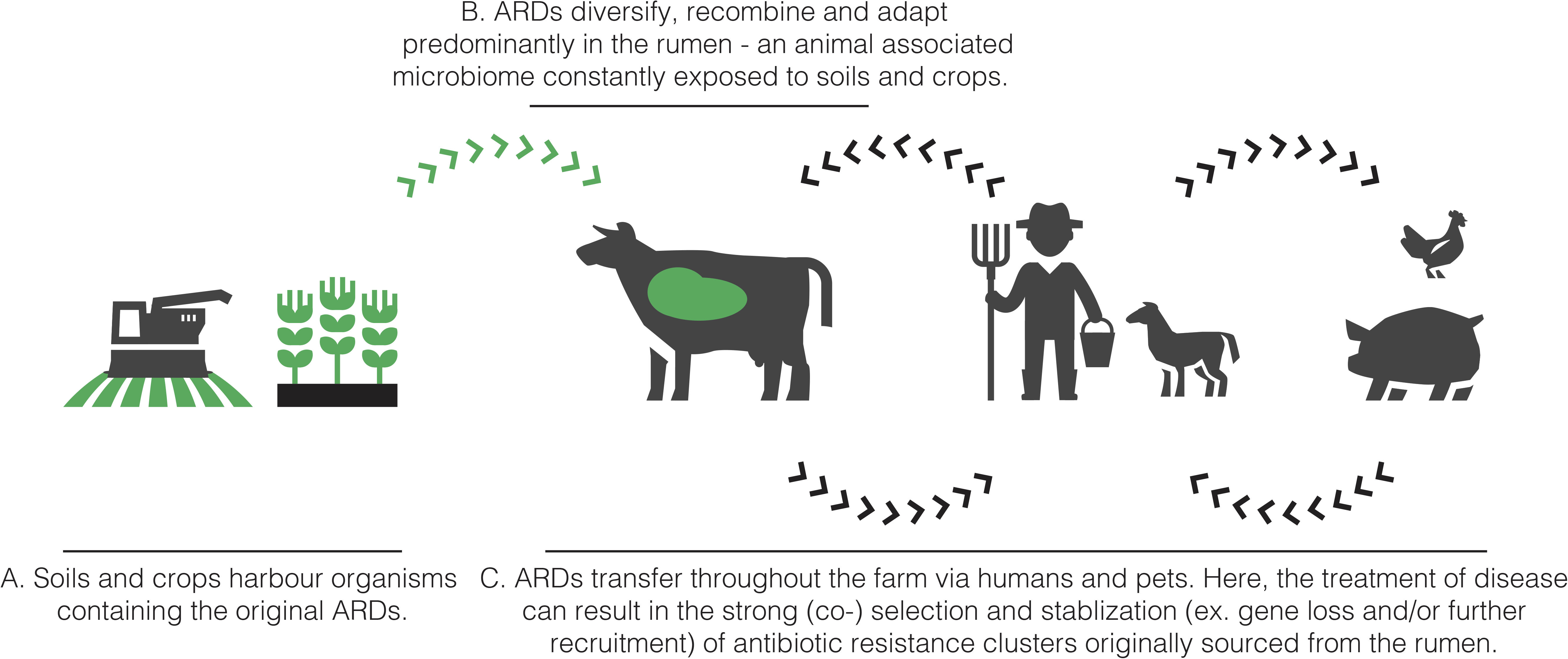
Graphical overview of a scenario where the rumen plays a predominant role in connecting soils and crops, harboring the organisms who produce antibiotics and have evolved ARDs, to the rest of the farm. It is suggested that the rumen provides a significant opportunity for ARDs to transfer into an animal associated microbiome, recombine and adapt before being spread throughout the farm, where the resulting antibiotic resistance cassettes are (co-) selected for in treating human or animal pathogens.

## METHODS

### Comparative analysis of *aadE-sat4-aphA-3* and *aadE-Ib*

Genome sequences from the Hungate1000 project, including those listed as previously published, were combined with MAGs from Steward *et al.*^15,16^. Using a concatenated fasta file containing all genome and MAG nucleotide sequences, ORFs were predicted using prodigal v2.6.3^49^. The resulting ORFs were then blasted against the CARD using local blastp v2.9.0+^50^. The 3 contigs (Hungate collection 4309689_79 and 43809680_52, MAG RUG782_1) which coded for the top 6 blast hits, in terms of bitscore, were then blasted against the NCBI nucleotide collection (nr/nt) using web-based blastn and the full-length sequence for each of the top 50 hits was downloaded^33,34^. After removing identical sequences, a total of 54 sequences were used in the downstream analysis, the accession numbers and descriptions for which are listed in Table S2. Each of the downloaded sequences was then blasted against the rumen sourced *Prevotella* sp. contig (RUG782_1) using local blastn 2.9.0+^50^. A sequence was displayed in Figure 1A if the total combined length of alignments was over 4000 bp for each query and the percent identity of the alignment was over 80%. For the gene diagrams displayed in Figure 1B, annotations from up to 10 of the top blastp hits from the NCBI non-redundant protein sequences database are listed in Table S3^33,34^. The same was done for Figure 3A, except starting with the contig from *B. thetaiotoamicron* nale-zl-c202 genome (43809680_59) containing *aadE-Ib*. Again, top 50 hits from the NCBI nucleotide collection (nr/nt) were downloaded (Table S2) and subsequently blasted against the rumen *B. thetaiotoamicron* contig (43809680_59). Here, a sequence was displayed in Figure 3A if the total combined length of alignments was over 1000 bp and the percent identity of the alignments were over 80%.

### Phylogenetic and syntenic analysis of AadE

The predicted ORF for AadE from the 3 selected rumen contigs (Hungate collection 4309689_79 and 43809680_52, MAG RUG782_1), being identical ORFs, was blasted against the NCBI non-redundant protein sequences database^33,34^. All hits with an e-value below 1e-4 were downloaded. Sequences were further eliminated if the length was below 250 bp or above 350 bp and an initial alignment was then made using MUSCLE (including the following flags:-maxiters 3 -diags -sv -distance1 kbit20_3)^51^. This alignment was inspected using Geneious v9.1.8, trimmed to between position 64 and 609, and further refined using the default setting from MUSCLE, while allowing for up to 50 iterations^51,52^. The phylogenetic tree shown in Figure 2 was subsequently constructed FastTree on the default settings^53^. In terms of visualization, clades were collapsed whose average branch length to the leaves was below 1.5 using the interactive tree of life (iTOL) online tool^54^. The resulting tree is down in Figure 2.

The tree shown in Figure 4A was constructed using the sequences extracted from clade 7 in Figure 2, with the addition of any homologues of AadE-Ia or AadE-Ib (>200 amino acids and >60% identity to the two versions from *B. thetaiotoamicron* nale-zl-c202 when compared using local blastp 2.9.0+) from the genomes used in Figure 1 and 3, if they contained multiple copies of the homologues^50^. The sequences were aligned using MUSCLE with 50 iterations, inspected using Geneious v9.1.8, and trimmed to between positions 25 and 305. Moreover, truncated proteins were removed, resulting in an alignment of 156 sequences, which was again refined using MUSCLE. This was then used as the input file for IQ-TREE using the standard settings with the following flags: -m TEST -bb 1000 -alrt 1000. An Le Gascuel (LG) model was selected using Gamma with 4 categories for the rate of heterogeneity^55–57^. The resulting tree, along with ultrafast bootstrap values, was visualized using iTOL.

To analyze synteny, any ORFs annotated as ARDs surrounding the AadE-Ia or AadE-Ib homologues (within maximum ~50kB) that were taken from the genomes used in Figure 1 and 3 are shown in Figure 4B and C. These were compared to AadE-Ib (Figure 4B) AadE-Ia (Figure 4C) or using local blastp 2.9.0+^50^. The annotation based on the top blastp hits from the NCBI non-redundant protein sequences database are listed in Table S5^33,34^.

## AKNOWLEDGMENTS

The competence centre FFoQSI is funded by the Austrian ministries BMVIT, BMDW and the Austrian provinces Niederoesterreich, Upper Austria and Vienna within the scope of COMET - Competence Centers for Excellent Technologies. The programme COMET is handled by the Austrian Research Promotion Agency FFG (Oesterreichische Forschungsförderungsgesellschaft). This work was conducted within the COMET-Project D4Dairy (Digitalisation, Data integration, Detection and Decision support in Dairying, Project number: 872039).

